# Apoptotic Cells Promote Tim4+ Macrophage Differentiation Through Direct and Indirect Signaling Pathways

**DOI:** 10.64898/2026.10.12.758552

**Authors:** Bryan S. Manning, Linna Zhu, Brennah Murphy, Marlaine M. Soliman, Chunlei Shao, Nesteene Param, Taito Miyamoto, Rahul Shinde, Martin Carroll, Nan Zhang

## Abstract

Tissue macrophages are the primary cells responsible for efficiently clearing apoptotic cells, yet how professional efferocyte populations are established remains unclear. Tim4 marks a subset of macrophages across multiple tissues that are highly efferocytic, but the mechanisms governing Tim4 acquisition are poorly understood. Here, we show that monocyte-derived macrophages progressively differentiate into Tim4⁺ macrophages in homeostatic and pathological settings. We identify a cell-autonomous pathway in which efficient apoptotic cell clearance promotes Tim4 acquisition in macrophages, as combined deficiency of CD11b and MerTK impairs both efferocytosis and Tim4 acquisition. Surprisingly, excessive apoptotic cell burden enhanced Tim4 acquisition independently of these receptors. We demonstrate that serial exposure to apoptotic cells induces a type 2 immune response characterized by expansion of IL-4-producing populations and that IL-4R-STAT6 signaling is sufficient and required to drive Tim4 acquisition. These findings support a model in which Tim4 acquisition in macrophages integrates direct signals associated with apoptotic cell clearance and indirect signals generated by apoptotic cell burden. We propose that these complementary mechanisms enable tissues to dynamically adjust the size and function of the professional efferocyte compartment to meet changing demands for apoptotic cell clearance.

## Introduction

The clearance of apoptotic cells, a process called efferocytosis, is a highly conserved mechanism among tissue macrophages. To efficiently and effectively clear apoptotic cells, and to prevent their decay towards secondary necrotic cells, macrophages express a wide repertoire of surface receptors to identify and engulf apoptotic cells^1–6^. Among these receptors is T-cell immunoglobin mucin-like domain 4 (Tim4) which directly binds to phosphatidylserine exposed on the surface of apoptotic cells and helps facilitate uptake^7–10^. Unlike several other efferocytosis receptors, Tim4 does not require a bridge molecule, like Gas6 or Protein S, for apoptotic cell recognition, enabling Tim4+ macrophages to rapidly tether and clear apoptotic targets even in dynamic tissue environments, such as the peritoneal fluid and the liver sinusoid.

Tim4 is expressed by specialized subsets of tissue-resident macrophages across multiple organs and has become widely used as a marker of time of residency^11–14^. In many tissues, Tim4 expression identifies macrophages with particularly robust efferocytic capacity, suggesting that it marks a professional efferocyte phenotype. While developmental and environmental cues that contribute to Tim4 expression have begun to emerge^14–19^, the mechanisms governing Tim4 acquisition remain poorly understood. In particular, it is unknown whether the process of efferocytosis itself contributes to the establishment of Tim4 expression.

Given the central role of Tim4 in apoptotic cell recognition, we hypothesized that interactions with apoptotic cells may provide instructive signals that drive Tim4 acquisition. Such a mechanism would allow tissue macrophages to adapt their efferocytic capacity to local demand, enabling macrophages that frequently encounter apoptotic cells to progressively acquire a more specialized efferocyte phenotype. Here, we demonstrate that Tim4 acquisition in macrophages is regulated by both cell-autonomous and cell non-autonomous mechanisms. We identify integrin αM (CD11b) as an important contributor to Tim4-independent efferocytosis and show that combined disruption of CD11b and the canonical efferocytosis receptor MerTK impairs Tim4 acquisition in macrophages at baseline and following bone marrow transplantation and adoptive transfer. In addition, we find that excessive apoptotic cell burden induces a type 2 immune response characterized by elevated IL-4 expression, which promotes Tim4 acquisition independently of macrophage efferocytic receptors. Together, these findings reveal that both apoptotic cell clearance and the immune responses generated by apoptotic cell accumulation contribute to the acquisition of the Tim4^+^ macrophage phenotype, providing new insight into how professional efferocyte populations are established and maintained.

## Results

### Monocyte-derived cells differentiate into Tim4+ macrophages in multiple settings

To investigate the mechanisms regulating Tim4 acquisition, we focused on macrophages within the peritoneal cavity, where both Tim4⁺ and Tim4⁻ macrophage populations coexist at steady state^15,16^. Consistent with previous reports^14,16^, analysis of 8-10 week-old C57BL/6J mice identified both populations (Fig. 1A-B), and the relative abundance of Tim4⁻ macrophages increased with age (Fig. 1C). Because Tim4 expression has historically been associated with embryonically derived macrophages, we first sought to determine whether monocyte-derived cells could acquire Tim4 expression within the peritoneal environment.

**Figure 1.**
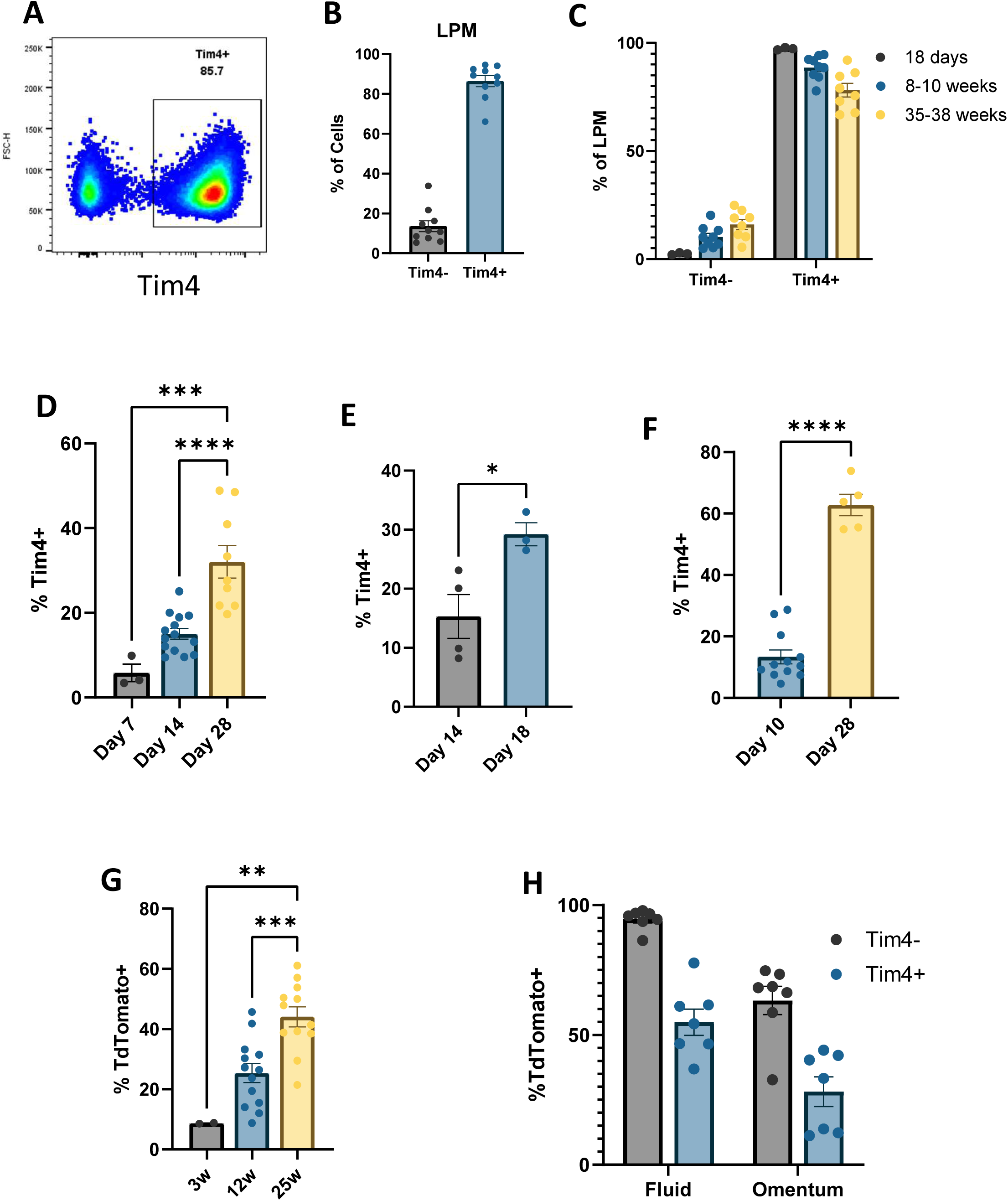
Monocytes differentiate into Tim4^+^ macrophages in multiple settings. (A) Representative dot plot of Tim4^+^ and Tim4^-^ peritoneal macrophages isolated from 10-week-old C57Bl6/J mice pre-gated on CD45^+^ ICAM2^Hi^ F4/80^Hi^ cells (B) Percentages of Tim4^+^ and Tim4^-^ peritoneal macrophages from 10-week-old mice. (C) Percentages of Tim4^+^ and Tim4^-^ peritoneal macrophages from 18-day old, 8-10-week-old, and 35-38-week-old mice (D) Percentage of donor CD45.1^+^ bone marrow monocytes that differentiated into Tim4^+^ peritoneal macrophages after the indicated time frame. ***p = 0.0001, ****p < 0.0001, one-way ANOVA with Tukey’s multiple comparisons test. (E) Percentage of donor CD45.1^+^ FACS-isolated Tim4^-^ peritoneal macrophages that differentiated into Tim4+ macrophages. p = 0.0305(*), student’s t-test. (F) Percentage of donor CD45.1^+^ FACS-isolated monocyte-macrophages from mice 72 hours after treatment with 1mg of Zymosan. ****p < 0.0001 student’s t-test. (G) Labeling kinetics TdTomato in Tim4^+^ peritoneal macrophages (CD45^+^ ICAM2^Hi^ F4/80^Hi^) from *Ms4a3^Cre^; Rosa26^LSL-TdT^* mice at different ages. **p = 0.0010, ***p = 0.0009, one-way ANOVA with Tukey’s multiple comparisons test. (H) TdTomato labeling in peritoneal and omental macrophages from *Ms4a3^Cre^; Rosa26^LSL-TdT^* mice 3 weeks after KPCA seeding.

To validate that monocytes-derived cells could in fact differentiate into Tim4^+^ cells, we employed adoptive transfer models to determine how readily monocyte and monocyte-derived cells can differentiate within the peritoneal cavity. Isolated CD45.1^+^ bone marrow monocytes adoptively transferred into the peritoneal cavity of CD45.2^+^ mice differentiated into peritoneal macrophages and steadily differentiated into Tim4^+^ cells over several weeks (Fig. 1D). In addition, adoptive transfer of sorted CD45.1^+^ Tim4^-^ peritoneal macrophages also confirmed that Tim4-macrophages can become Tim4^+^ over time in CD45.2^+^ mice (Fig. 1E). To model this differentiation during the resolution from sterile inflammation, inflammatory monocyte-derived macrophages sorted from zymosan treated CD45.1^+^ mice were adoptively transferred into CD45.2^+^ mice that were treated with an equivalent dose of zymosan on the same day to mimic their native environment^20^. Here too, adoptively transferred cells differentiated into Tim4^+^ macrophages over several weeks (Fig. 1F). Together, these complementary adoptive transfer models demonstrate that monocytes and monocyte-derived macrophages can progressively acquire Tim4 expression within the peritoneal cavity.

Although adoptive transfer studies demonstrated that monocyte-derived cells can acquire Tim4 expression, these experimental systems do not address the extent to which monocytes contribute to the Tim4⁺ macrophage pool under physiological conditions. Lineage tracing models have determined that the Tim4^-^ cells are entirely monocyte derived cells^21^. Historically, Tim4⁺ peritoneal macrophages were considered a predominantly embryonically derived population with minimal contribution from circulating monocytes^22^. However, recent lineage-tracing studies have revealed substantially greater monocyte input into tissue macrophage populations than previously appreciated^21^. Employing the *Ms4a3^Cre^; Rosa26^LSL-TdT^* mouse model, we determined that as mice age, the contribution of monocyte derived cells to the peritoneal Tim4^+^ population steadily increases (Fig. 1G). We additionally used the *Ms4a3^Cre^; Rosa26^LSL-TdT^* lineage tracing mouse to demonstrate that during metastatic ovarian cancer, monocyte derived cells contribute to Tim4^+^ macrophages in both the fluid and omental tumor compartments (Fig 1H). Collectively, these data demonstrate that Tim4 expression is not restricted to embryonically derived macrophages and establish that monocyte-derived cells contribute substantially to the Tim4⁺ macrophage compartment during both homeostasis and disease. These findings indicate that Tim4 expression can be acquired after tissue entry and therefore is unlikely to be determined solely by developmental origin. Instead, the progressive acquisition of Tim4 by monocyte-derived cells suggests that signals within the tissue environment regulate the establishment of the Tim4⁺ macrophage phenotype.

### Tim4^-^ macrophage efferocytosis is dependent on integrin α_M_β_2_

Having established that monocyte-derived macrophages can acquire Tim4 expression after entering tissues, we next sought to identify the environmental signals that regulate this process. One clear functional distinction between Tim4^+^ and Tim4^-^ macrophages is their efficiency of performing efferocytosis. Consistent with previous reports demonstrating a role for Tim4 in apoptotic cell tethering and engulfment^8,10,12,18^, we found that Tim4⁺ macrophages cleared apoptotic cells more efficiently than their Tim4⁻ counterparts, although Tim4⁻ macrophages retained substantial efferocytic activity (Fig. S1A-B). These observations suggested that Tim4 expression may mark a specialized efferocyte subset rather than simply a developmentally distinct macrophage population. We therefore hypothesized that the process of efferocytosis itself provides instructive signals that promote Tim4 acquisition. Such a mechanism would establish a feed-forward circuit in which macrophages that successfully engage apoptotic cells progressively acquire enhanced efferocytic capabilities through Tim4 expression. This model could also explain previous observations that mature resident macrophages suppress Tim4 differentiation from incoming cells^12,21^, as established Tim4⁺ macrophages would be expected to outcompete immature Tim4⁻ macrophages for access to apoptotic cell cargo. To test this hypothesis, we sought to identify the receptors responsible for efferocytosis by Tim4⁻ macrophages and determine whether disruption of these pathways altered Tim4 differentiation. We first checked MerTK as a canonical efferocytosis receptor. Bone marrow derived macrophages (BMDMs) generated from *Mertk^-/-^* mice demonstrated an efferocytosis deficiency relative to wildtype (WT) cells when fed apoptotic cells in vitro (Fig S1C-D), but surprisingly we found little difference between WT and *Mertk^-/-^* Tim4^-^ macrophages when apoptotic cells were injected intraperitoneally (Fig S1E-G). Conversely, we found that *Mertk^-/-^* Tim4^+^ peritoneal macrophages demonstrated slightly decreased efferocytosis capacity relative to WT macrophages (Fig S1H-J), supporting the finding that MerTK supports Tim4-driven phagocytosis as shown by others^23,24^. Because MerTK deficiency had surprisingly little impact on the efferocytosis by Tim4⁻ peritoneal macrophages, we reasoned that another receptor must support apoptotic cell clearance in this population. One notable candidate was CD11b (integrin α_M_, *Itgam*), which is expressed at exceptionally high levels on peritoneal macrophages despite their persistence within a fluid environment^25,26^. Together with CD18 (integrin β_2_), CD11b forms the heterodimer integrin α_M_β_2_ also known as Mac-1 or complement receptor 3 (CR3). Because CR3 has previously been reported to recognize complement-opsonized apoptotic cells^27–30^, and because peritoneal macrophages express exceptionally high levels of CD11b, we investigated whether this pathway contributes to efferocytosis in Tim4^⁻^ macrophages. BMDMs generated from *Itgam^-/-^* mice demonstrated decreased efferocytosis in vitro, although to a lesser extent than *Mertk^-/-^* BMDMs (Fig S1C-D). Surprisingly, intraperitoneal injections of apoptotic cells into WT and *Itgam^-/-^* mice revealed that Tim4^-^ macrophages exhibited drastic deficiency in apoptotic cell clearance in *Itgam^-/-^* mice (Fig S1E-G), while Tim4^+^ macrophages showed little change (Fig S1H-J), suggesting a critical but underappreciated role of CD11b in apoptotic cell clearance in vivo. We also validated this finding in an integrin signaling deficient mouse model, *Lyz2*^Cre^; *Tln1*^fl/fl^ (Fig S1K). Thus, CD11b but not MerTK seems to be integral for efficient efferocytosis by Tim4^-^ peritoneal macrophages.

### Double Knockout of CD11b and MerTK demonstrates Tim4^+^ differentiation impairment

To investigate whether impaired efferocytosis affected Tim4^+^ differentiation, we evaluated several models of monocyte and monocyte-derived macrophage transfer using *Itgam*⁻^/^⁻ cells. In a competitive bone marrow transplantation model, WT and *Itgam*⁻^/^⁻ donor cells generated comparable frequencies of Tim4⁺ macrophages, indicating a dispensable role of CD11b in Tim4 acquisition in macrophages (Fig. S2A). We next adoptively transferred magnetic-activated cell sorted (MACS) purified bone marrow monocytes from WT (CD45.1) and *Itgam*⁻^/^⁻ (CD45.2) mice into congenic recipients and assessed Tim4 acquisition over time. Although female mice exhibited genotype-dependent differences, with *Itgam*⁻^/^⁻ monocytes showing reduced Tim4 expression at 2 weeks but increased Tim4 expression at 4 weeks, these effects were inconsistent across experimental settings (Fig. S2B-C). Similarly, following sterile inflammation induced by zymosan, adoptively transferred *Itgam*⁻^/^⁻ inflammatory mono-macrophages displayed modest defects in Tim4 acquisition that were again restricted to female mice (Fig. S2D-E). Collectively, these findings indicate that although CD11b contributes substantially to efferocytosis by Tim4⁻ macrophages, loss of CD11b alone has only limited and inconsistent effects on Tim4 acquisition in monocyte-derived macrophages, suggesting that compensatory efferocytosis pathways preserve their capacity of Tim4 acquisition.

Given the substantial redundancy among efferocytosis receptors, we reasoned that deletion of a single receptor may be insufficient to disrupt Tim4 acquisition. CD11b and MerTK have previously been shown to function cooperatively during efferocytosis^31,32^ suggesting that concurrent disruption of both pathways might overcome compensatory mechanisms that preserve apoptotic cell clearance. To test this possibility, we generated *Itgam*^−/−^;*Mertk*^−/−^ double-knockout (DKO) mice. We initially generated the DKO by crossing the *Itgam^-/-^* strain with the long studied *Mertk^-/-V1^* strain. Analysis of DKO mice at homeostasis revealed an increased frequency of Tim4⁻ peritoneal macrophages (Fig 2A-B), which resulted from an expansion in the absolute number of Tim4⁻ macrophages (Fig 2C-D). To directly assess Tim4 acquisition, we performed competitive bone marrow transplantation using WT CD45.1 and DKO CD45.2 donor marrow transplanted into lethally irradiated CD45.2 recipient mice (Fig. 2E). In contrast to the modest phenotype observed with CD11b deficiency alone, DKO cells demonstrated a consistent defect in Tim4 acquisition throughout immune reconstitution at 2, 4, and 6 weeks post-transplantation (Fig. 2F-H). Interestingly, this phenotype was only apparent under competitive conditions. In a non-competitive reconstitution model, DKO macrophages exhibited no detectable defect in Tim4 acquisition (Fig. S3A), suggesting that loss of CD11b and MerTK reduces the competitive fitness of macrophages during establishment of the Tim4⁺ compartment rather than completely preventing Tim4 acquisition. These findings support a model in which competition for apoptotic cell cargo influences Tim4 acquisition, consistent with previous reports demonstrating that resident macrophages can restrict the maturation of incoming monocyte-derived cells into Tim4⁺ macrophages^20^. To confirm that our DKO cells had defective efferocytosis, we injected apoptotic cells into these mice and found clearance defects in both Tim4^+^ and Tim4^-^ DKO peritoneal macrophages relative to WT cells (Fig 2I-K).

**Figure 2.**
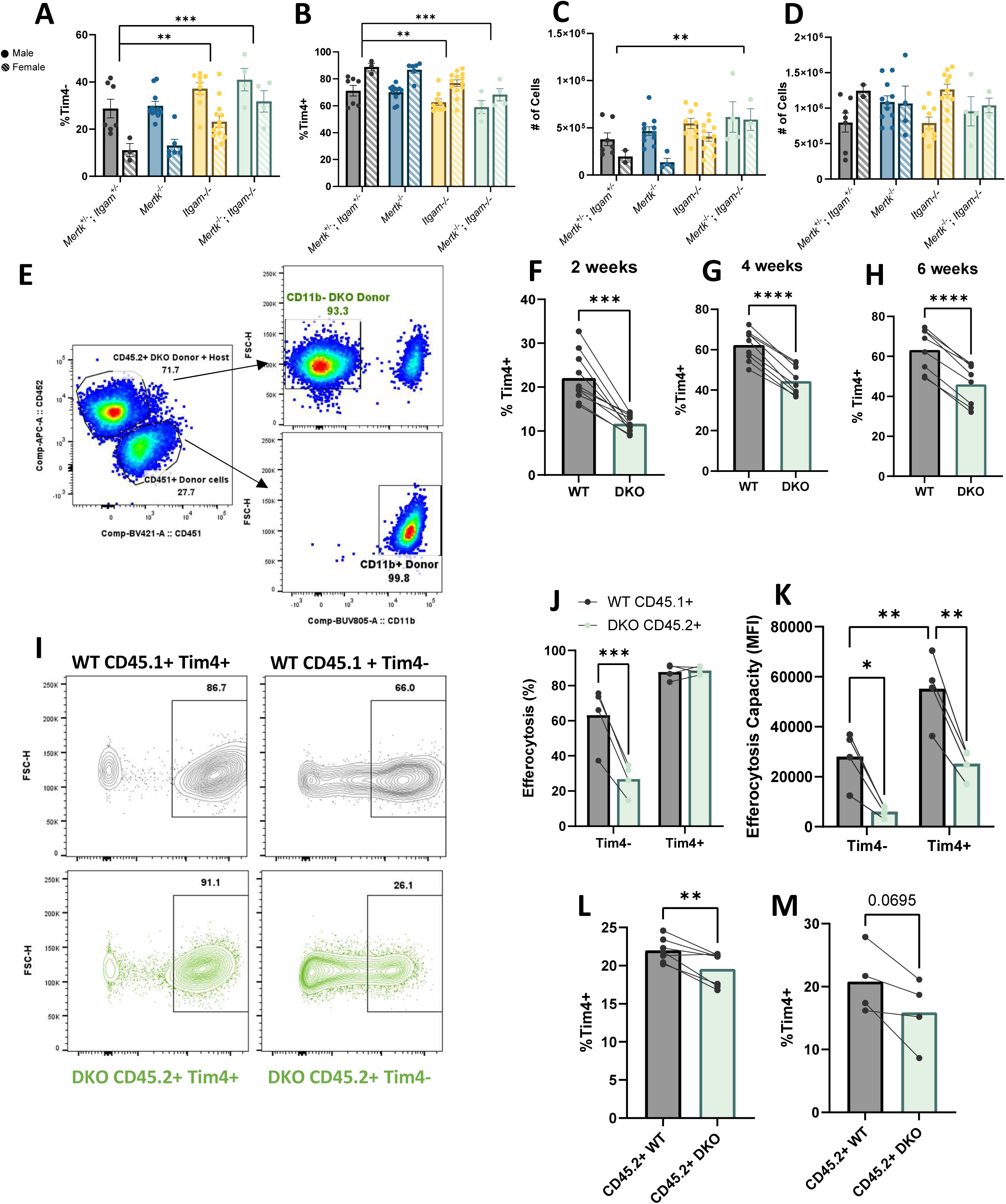
Double knockout of both integrin αMβ2 and MerTK results in defective efferocytosis and increased Tim4^-^ cells. (A-B) Proportion and (C-D) Quantification of peritoneal macrophages that are Tim4^-^ and Tim4^+^ from Het;Het, Mertk^-/-^, Itgam^-/-^, and DKO 8–10-week-old mice. Data are presented as mean ± SEM. Statistical analysis was performed using two-way ANOVA with Tukey’s multiple-comparisons test. P values shown indicate significant differences for the main effect of genotype (column factor), independent of sex. *P < 0.05, **P < 0.01, ***P < 0.001, ****P < 0.0001. (E) Flow cytometry for the discrimination of donor CD45.1^+^ CD11b^Hi^ WT and CD45.2^+^, CD11b^-^ DKO and host CD45.2^+^, CD11b^Hi^ peritoneal macrophages from mixed bone marrow chimeras. (F-H) Percentages of Tim4^+^ from CD45.1 WT and CD45.2 DKO peritoneal macrophages from mixed bone marrow chimeras (pooled from two experiments per timepoint). Each point represents an individual mouse. Lines connect paired measurements from the same mouse. Statistical significance was assessed using a paired two-tailed Student’s t-test. *P < 0.05, **P < 0.01. (I) DiD-labeled apoptotic Jurkat cells were injected i.p. in mixed bone marrow chimera mice 6 week after transplant and peritoneal macrophages were analyzed for AC uptake after 1 hour. Both Tim4+ and Tim4-macrophages from both donors were gated on AC+. (J-K) Efferocytosis efficiency (DiD+ macrophages) and capacity (DiD mean fluorescence intensity, MFI) in macrophages. (L) Percentages of Tim4^+^ CD45.2 WT and CD45.2 DKO peritoneal macrophages18 days after adoptive transfer of BMDMs into NSG mice. (M) Percentages of Tim4^+^ CD45.2 WT and CD45.2 DKO 18 days after adoptive transfer of bone marrow monocytes into NSG mice.

We next sought to rule out alternative explanations such as decreased trafficking or increased cell death of Tim4^+^ macrophages. While CD11b has known roles in immune cell trafficking, competitive BMT mice revealed an increase of DKO chimerism relative to WT (Fig S3B-D) at each time point. Both *in vitro* and *in vivo* analysis of WT and DKO peritoneal macrophages revealed that DKO peritoneal macrophages seemingly had a survival advantage relative to WT cells, perhaps due to loss of CD11b as has been previously reported^33^. We additionally performed adoptive transfer experiments with CD45.2 WT and CD45.2 DKO cells into NSG mice and observed that DKO BMDMs and monocytes had Tim4 acquisition defects (Fig 2L-M). Because recent reports have suggested that phenotypes observed in the original *Mertk^−/−V1^* strain may be influenced by additional genetic alterations^34^, we sought to validate our findings using the independently generated *Mertk^−/−V2^* allele. V2-derived DKO mice retained Tim4 differentiation defects both at homeostasis (Fig S3E-H) and in our BMT model (Fig S3I-J). Although the phenotype observed in the V2-derived DKO was less pronounced than that observed with the V1-derived DKO, the persistence of a significant Tim4 differentiation defect across both strains suggests that CD11b and MerTK contribute to Tim4 acquisition while also highlighting the likelihood that additional compensatory pathways participate in this process. Collectively, these findings indicate that simultaneous disruption of CD11b- and MerTK-mediated efferocytosis impairs acquisition of the Tim4⁺ macrophage phenotype. The requirement for combined receptor deficiency and the preferential defect observed under competitive conditions suggest that efficient apoptotic cell clearance provides a selective advantage during establishment of the Tim4⁺ macrophage compartment.

### Exogenous apoptotic cells drive Tim4^+^ differentiation independent of efferocytosis ability

Having found that disruption of CD11b- and MerTK-mediated efferocytosis impaired Tim4 differentiation, we next asked whether increasing access to apoptotic cells could accelerate acquisition of the Tim4⁺ phenotype. To test this possibility, BMDMs were adoptively transferred into the peritoneal cavity and recipient mice were serially administered apoptotic thymocytes every few days throughout the differentiation period (Fig. 3A). Following five apoptotic cell injections over 18 days, transferred macrophages exhibited increased Tim4 differentiation compared to PBS-treated controls (Fig. 3B-C). This effect was reproducible when host macrophages were depleted with clodronate-loaded liposomes prior to transfer (Fig. 3D) and when Tim4⁻ peritoneal macrophages were used as donor cells (Fig. 3E), indicating that enhanced Tim4 acquisition was not restricted to a particular donor population or niche occupancy state. In addition to donor-derived cells, endogenous host macrophages also exhibited increased Tim4 expression following apoptotic cell administration (Fig. 3F-G). We next asked whether this increase in Tim4 acquisition required CD11b- and MerTK-mediated efferocytosis. Unexpectedly, apoptotic cell administration enhanced Tim4^+^ differentiation to a similar extent in both WT and efferocytosis-deficient macrophages (Fig. 3H-I). These findings suggested that excess apoptotic cells promote Tim4 acquisition through a mechanism distinct from the cell-autonomous efferocytosis pathways identified above. We therefore hypothesized that apoptotic cell burden alters the peritoneal immune environment in a manner that indirectly promotes Tim4^+^ differentiation.

**Figure 3.**
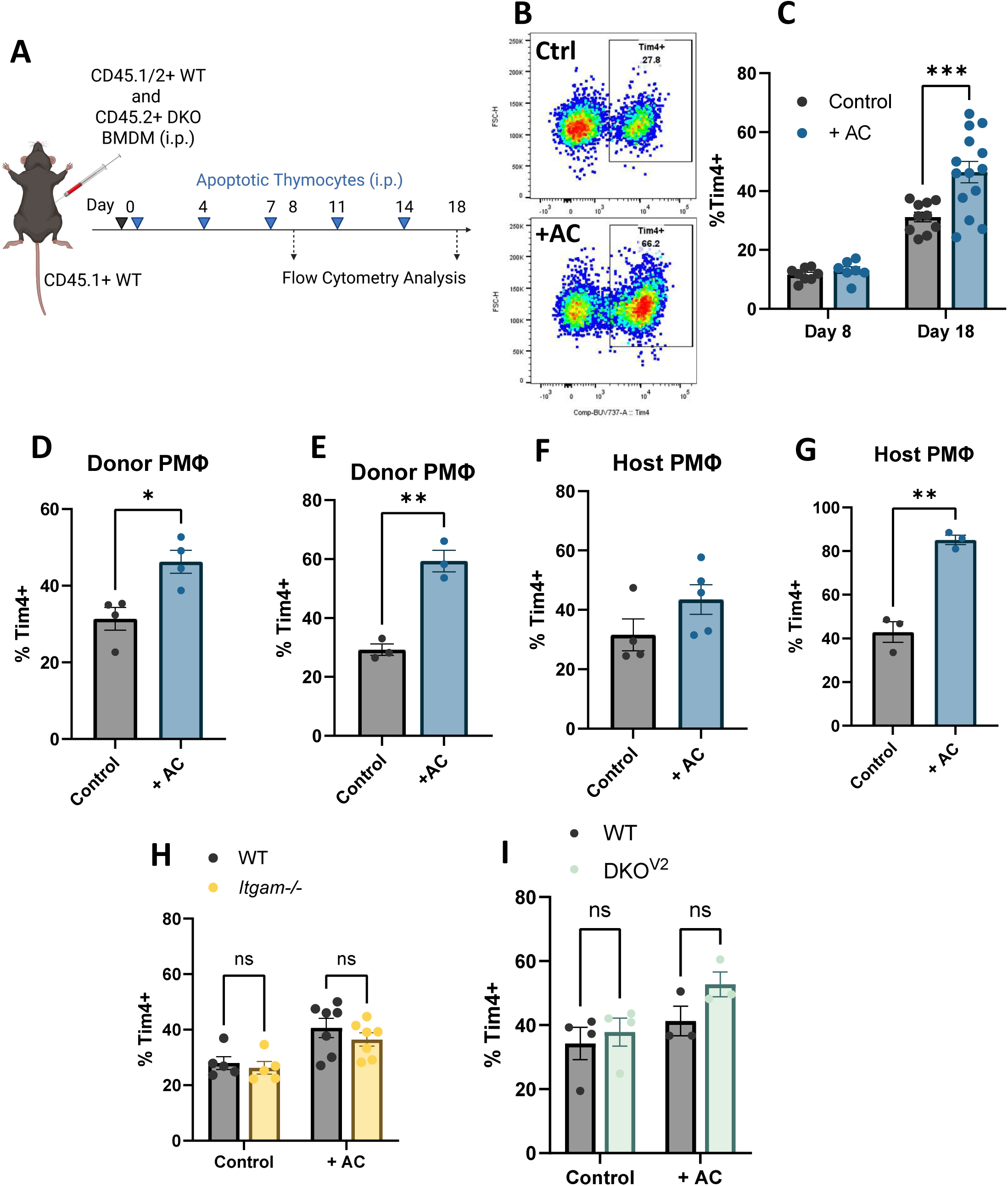
Exogenous apoptotic cells accelerate Tim4^+^ differentiation. (A) Experimental scheme for transfer of BMDMs and apoptotic thymocytes into recipient mice. (B) Representative dot plot of Tim4^+^ and Tim4^-^ peritoneal macrophages from control mice and serial AC-injected mice at day 18. (C) Proportion of Tim4^+^ donor peritoneal macrophages 8 and 18 days after BMDM adoptive transfer. ***p = 0.0003, two-way ANOVA with Tukey’s multiple comparisons test. (D) Proportion of Tim4^+^ donor peritoneal macrophages 18 days after BMDM adoptive transfer into clodronate-depleted recipients (E) Proportion of Tim4^+^ donor peritoneal macrophages 18 days after Tim4^-^ peritoneal macrophage adoptive transfer. (F-G) Proportion of Tim4^+^ host peritoneal macrophages from experiments in (D) and (E) respectively (H-I) Proportion of Tim4^+^ donor peritoneal macrophages 18 days after competitive adoptive transfer of CD45.2^+^ CD11b^Hi^ WT and CD45.2^+^ CD11b^-^ Itgam KO or DKO +/- apoptotic cell injections.

### Exogenous apoptotic cells spark type 2 immune response and drives IL-4-STAT6-dependent Tim4 acquisition

Because apoptotic cell administration enhanced Tim4^+^ macrophage differentiation independently of CD11b and MerTK, we next sought to determine whether excess apoptotic cells altered the peritoneal immune environment in a manner that indirectly promoted Tim4 acquisition. Serial administration of apoptotic cells resulted in a marked increase in overall peritoneal cellularity (Fig. 4A), characterized by expansion of host macrophages, T cells, and B cells, as well as a prominent influx of eosinophils (Fig. 4B). Together, these changes resembled a type 2 immune response^35–38^. To assess production of the canonical type 2 cytokines IL-4 and IL-13, peritoneal exudate cells were restimulated with anti-CD3/CD28 prior to intracellular cytokine staining. While the frequency of IL-4-producing CD4^+^ T cells was similar between control and apoptotic cell-treated mice (Fig. S4A), the expansion of the CD4^+^ T-cell compartment resulted in an increased number of IL-4-producing T cells within the peritoneal cavity. In addition, both host peritoneal macrophages and infiltrating eosinophils exhibited increased intracellular IL-4 staining following apoptotic cell administration (Fig. 4C-D). Intracellular IL-13 levels were more variable in CD4+ T cells, eosinophils and macrophages (Sup Fig 4B-D). Given the established role of IL-4 in shaping macrophage responses, we next investigated whether IL-4 itself could promote Tim4^+^ macrophage differentiation.

**Figure 4.**
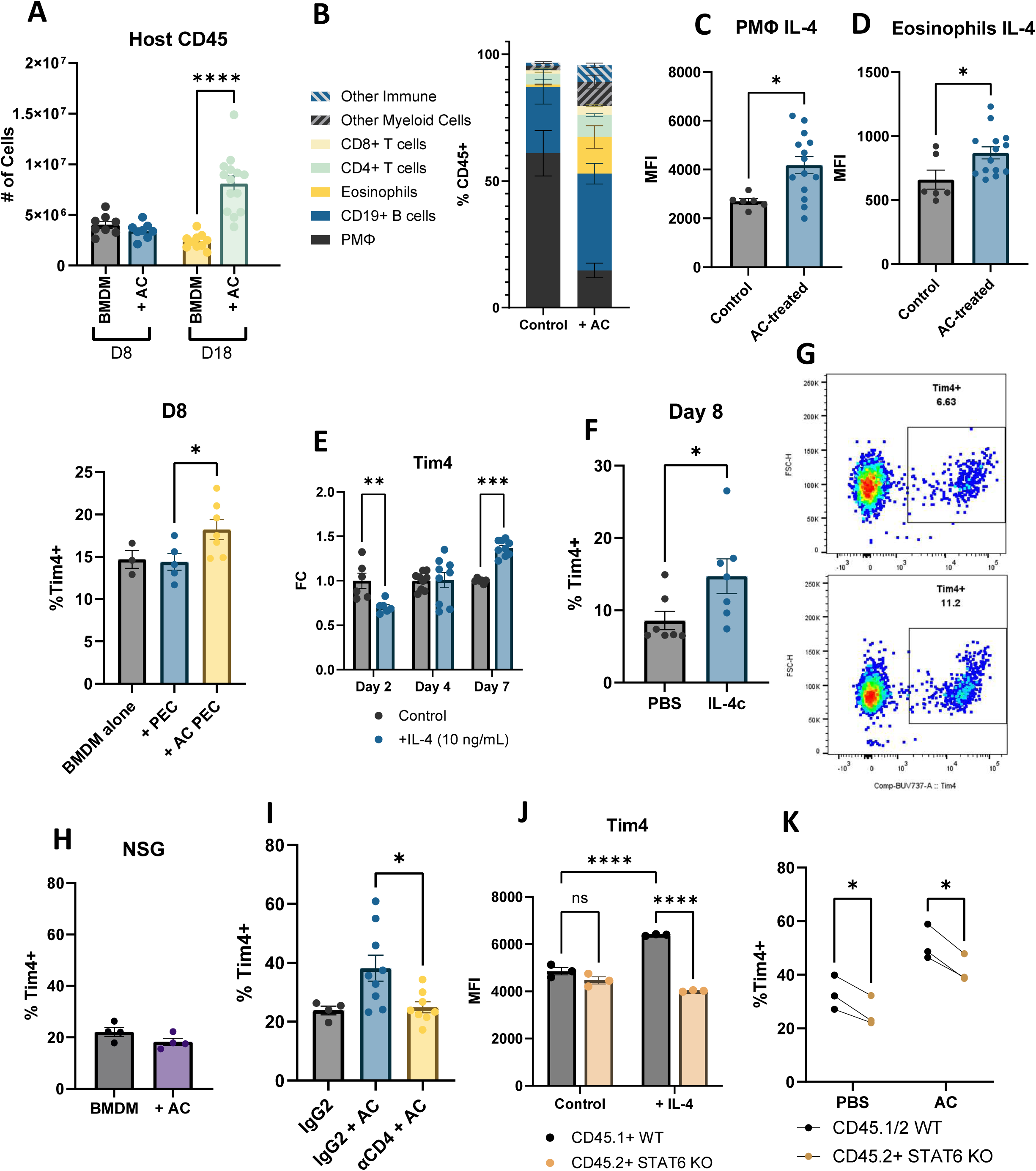
Exogeneous apoptotic cells generate type 2 response driving Tim4+ differentiation through IL-4R-STAT6 axis. (A) Quantification of total host CD45+ immune cells 8 and 18 days after adoptive transfer with serial apoptotic cell injections. ****p < 0.0001, two-way ANOVA with Tukey’s multiple comparisons test. (B) Composition of the host CD45+ population in the peritoneal fluid. (C-D) Intracellular IL-4 MFI of peritoneal macrophages and eosinophils on day 15 of the immune response. p < 0.05(*), student’s t-test. (E) Fold change of surface Tim4 expression on BMDMs after sustained treatment with recombinant IL-4 (10 ng/mL) after 2-, 4-, and 7-days. ** p < 0.01, *** p < 0.001, two-way ANOVA with Tukey’s multiple comparisons test. (F-G) Proportion of Tim4+ donor peritoneal macrophages 8 days after BMDM adoptive transfer in mice. Mice received IL-4 complex on days 1 and 4 after adoptive transfer. p < 0.05(*), student’s t-test (H) Proportion of Tim4+ donor peritoneal macrophages 18 days after BMDM adoptive transfer into NSG mice +/- apoptotic cell injections. (I) Proportion of Tim4+ donor peritoneal macrophages 18 days after BMDM adoptive transfer with isotype control or anti-CD4 treatment (J) WT and *Stat6^-/-^* BMDM Tim4 expression with or without 7 days of IL-4 (10 ng/mL). ****p < 0.0001, two-way ANOVA with Tukey’s multiple comparisons test. (K) Proportion of Tim4+ donor peritoneal macrophages 18 days after competitive adoptive transfer of CD45.1/2 WT and CD45.2 *Stat6^-/-^* +/- apoptotic cell injections.

To determine whether IL-4 was sufficient to promote Tim4 expression in vitro, BMDMs were cultured in the presence of IL-4. We observed a decrease in Tim4 expression after 2 days, however after a week of sustained treatment, we observed increases in Tim4 MFI (Fig 4E). Next, we performed an adoptive transfer of BMDMs into mice injected with two doses of IL-4 complex (IL-4c) and revealed increased donor Tim4 acquisition on day 8 (Fig 4F-G). To confirm that adaptive immune cells were required for apoptotic cell-driven Tim4 acquisition, we repeated these experiments in NSG mice and in mice depleted of CD4^+^ T cells. In both settings, apoptotic cell administration failed to enhance Tim4^+^ differentiation (Fig. 4H-I), demonstrating that adaptive immune populations are required for this response. Similar findings were observed in TCRα-deficient mice (Fig. S4E).

Because IL-4 signals primarily through STAT6 to drive alternative macrophage activation^39–41^, we next asked whether STAT6 was required for Tim4 induction. IL-4-mediated upregulation of Tim4 in cultured BMDMs was abolished in *Stat6^−/−^* BMDMs (Fig. 4J). Likewise, when WT and *Stat6^−/−^* BMDMs were adoptively transferred into mice receiving serial apoptotic cell injections, STAT6 deficiency impaired both baseline and apoptotic cell-induced Tim4 acquisition (Fig. 4K). These findings identify STAT6 as a critical downstream mediator linking IL-4 signaling to Tim4 acquisition. Collectively, these findings indicate that excessive apoptotic cell burden induces a type 2 immune response that promotes Tim4^+^ macrophage differentiation through a mechanism distinct from direct efferocytosis signaling. Rather than requiring CD11b- and MerTK-mediated apoptotic cell uptake, this pathway depends upon adaptive immune responses, IL-4 production, and STAT6 signaling within macrophages.

## Discussion

Our findings suggest that Tim4^+^ macrophage differentiation is driven in part through cell autonomous efferocytosis signaling in homeostatic conditions and cell non-autonomous IL-4R-STAT6 signaling in the presence of excessive numbers of apoptotic cells. These results provide a conceptual framework for how Tim4 expression increases with macrophage time-of-residency in a tissue niche and suggest that the integration of apoptotic cell sensing and local immune cues shapes the professional efferocyte phenotype. Importantly, our adoptive transfer and lineage tracing studies demonstrate that Tim4 expression can be acquired by monocyte-derived macrophages in both homeostatic and pathological settings, indicating that Tim4 is not solely a consequence of developmental origin but rather a regulated macrophage state responsive to environmental cues. Numerous studies have identified tissue-specific factors that instruct resident macrophage identity, including retinoic acid-dependent regulation of GATA6 in peritoneal macrophages^22,42^ and DLL4-mediated induction of LXRa expression in Kupffer cells^43^. While these niche-derived signals generate tissue-specific specialization, a shared feature of resident macrophages is their capacity for efferocytosis. Tim4 has long been recognized as a marker of macrophage residency, yet the factors governing its acquisition have remained poorly understood. We hypothesized that sustained exposure to apoptotic cells might constitute one such signal. Consistent with the idea, disrupting efferocytosis by targeting CD11b and MerTK impaired Tim4 acquisition in peritoneal macrophages. Together, these findings support a feed-forward model in which engagement in apoptotic cell clearance promotes acquisition of the Tim4⁺ state, thereby generating increasingly specialized professional efferocytes.

Unexpectedly, serial administration of exogenous apoptotic cells increased Tim4^+^ macrophage differentiation even in contexts where efferocytosis signaling was severely impaired. This finding suggested that excessive apoptotic cells could promote Tim4 acquisition through a mechanism distinct from direct efferocytosis-associated signaling. Successive injections of apoptotic cells induced marked changes in the peritoneal immune landscape, including eosinophilia, lymphocyte expansion, and increased numbers of IL-4 producing cells, ultimately leading us to investigate cytokine-mediated pathways. IL-4 is a well-established regulator of tissue repair and macrophage activation, and previous studies have demonstrated that IL-4 can amplify efferocytosis-associated transcriptional programs and promote anti-inflammatory responses following apoptotic cell clearance^44,45^. Recent work has also characterized how IL-4 can modulate the macrophage pool by contextually increasing or decreasing monocyte and monocyte-derived macrophages contribution^46,47^. Our data build off these observations and further characterizes how IL-4 can promote the efferocytosis ability of monocyte-derived cells. While the dominant source of IL-4 within this response remains to be determined, our data demonstrate that IL-4 is sufficient to promote Tim4^+^ macrophage differentiation and that STAT6 is required for both IL-4-driven and apoptotic cell-driven Tim4 acquisition.

While the mechanisms that initiate type 2 immunity remain incompletely defined, our findings raise the possibility that apoptotic cell accumulation beyond normal homeostatic clearance capacity may contribute to the generation of type 2 responses. In this model, excessive cell death resulting from tissue injury, parasite infection, environmental insult, or cancer could create conditions that favor IL-4 production and encourage subsequent expansion of Tim4^+^ macrophages. In cancer particularly, the increased presence of Tim4+ macrophages and IL-4 signaling have both been shown to promote tumor and impair treatment response^48–51^. Our observations linking apoptotic cell accumulation to IL-4 production and Tim4 acquisition raise the possibility of a common pathway connecting apoptotic cell burden with the established protumor effects of IL-4 signaling and Tim4⁺ tumor-associated macrophages. Future studies will be needed to determine the mechanisms that mediate this apoptotic cell-induced type 2 immunity.

Although our results support a model in which efferocytosis-associated signaling and IL-4R-STAT6 signaling contribute to Tim4^+^ macrophage differentiation in the peritoneal cavity, the extent to which these mechanisms operate in other tissue macrophage populations remains unclear. Tim4^+^ macrophages are present in numerous tissues^11^, each characterized by distinct developmental origins, environmental cues, and apoptotic cell interactions. Future studies will be required to determine whether the pathways identified here represent a general mechanism governing Tim4 acquisition or a specialization of the peritoneal niche. While the requirement for CD11b and MerTK implicates efferocytosis in Tim4^+^ differentiation, the present studies do not fully distinguish between signaling events downstream of these receptors and the broader consequences of impaired apoptotic cell clearance. Reduced Tim4^+^ macrophage differentiation may reflect loss of receptor-specific signaling, diminished engulfment efficiency, altered processing of apoptotic cargo, or a combination of these effects. Dissecting these possibilities will require approaches that separate engulfment from receptor-mediated signal transduction.

More broadly, these observations raise the possibility that apoptotic cell burden serves not only as a homeostatic challenge requiring clearance, but also as an instructive signal that shapes macrophage identity and local immune responses. Determining how signals derived from engulfment are integrated with tissue-specific niche factors and cytokine cues will be an important goal for future studies. Such work may reveal whether the pathways identified here represent a conserved mechanism through which tissues adapt their efferocytic capacity to changing physiological demands.

In summary, we demonstrate that Tim4^+^ macrophage differentiation is regulated by both direct efferocytosis-associated signals and indirect IL-4-driven immune signals. Our findings establish that monocyte-derived macrophages can acquire the Tim4⁺ professional efferocyte phenotype and identify both apoptotic cell clearance and apoptotic cell burden as key regulators of this process. We propose that these complementary pathways allow tissues to adapt the size and function of their professional efferocyte compartment in response to changing demands for apoptotic cell clearance. By defining mechanisms governing Tim4 acquisition, this work advances our understanding of how resident macrophages develop specialized functions required to maintain tissue homeostasis during both steady-state turnover and periods of increased cellular stress.

## Methods

### Primary cell culture

To culture bone marrow-derived macrophages (BMDMs), bone marrow was isolated from 8–12-week-old mice and cultured for 5 days in DMEM supplemented with 10% heat-inactivated PBS, 10 U/mL penicillin, 100 mg/mL streptomycin and 40 ng/mL of recombinant murine CSF1 (BioLegend). After 5 days, cells were removed from petri dishes using pre-warmed Cell Stripper (Invitrogen) for 20-25 minutes at room temperature.

For longer term culture experiments, BMDMs were grown in DMEM supplemented with 10% heat-inactivated PBS, 10 U/mL penicillin, 100 mg/mL streptomycin, 20 ng/mL of recombinant CSF1 and if indicated 10 ng/mL of recombinant IL-4. Fresh media was exchanged on days 2 and 4.

### Animal models

C57BL/6J (WT, #000664), *Itgam^-/-^* B6.129S4-Itgamtm1Myd/J, #003991), Lyz2^Cre^ (B6.129P2-Lyz2tm1(cre)Ifo/J, #004781), *Rosa26^LSL-TdT^* (B6.Cg-Gt(ROSA)26Sortm14(CAG-tdTomato)Hze/J, # 007914), *Mertk^-/-v1^* (B6;129-Mertktm1Grl/J, # 011122), *Mertk^-/-v2^* (B6(129)-Mertktm1.2Cvro/J, #038128), Tcra^-/-^ (B6.129S2-Tcratm1Mom/J, # 002116), *Ms4a3^Cre^* (C57BL/6J-Ms4a3em2(cre)Fgnx/J, #036382), *Stat6^-/-^* (B6.129S2(C)-Stat6tm1Gru/J, #005977) mice were purchased from Jackson Laboratories. NSG mice were purchased from the Wistar Animal Facility. *Tln1^fl/fl^* mice were a kind gift from Dr. Rodger P. McEver (Oklahoma Medical Research Foundation, Oklahoma City, OK)

All mice were housed in individual microisolator cages in a rack system with filtered air in Wistar’s mouse barrier facility and provided with shelter and enrichment to reduce stress. Mice were kept on a 12-h light-dark cycle and had access to food and water ad libitum. All animal procedures were performed in accordance with the Wistar Institutional Animal Care and Use Committee under protocol 201536. Genotyping was performed utilizing Transnetyx automated genotyping services.

### Bone Marrow Transplant

Six- to ten-week-old male and female mice were irradiated using 10 Gy ionizing whole body radiation using SARRP. The same day, bone marrow cells were harvested from sex-matched donor mice of the indicated genotype. Animals were subsequently given 5 x 10^6^ BM cells per genotype by intravenous injection.

### Cell Isolation

Mice were sacrificed by exposure to rising levels of CO2. Peritoneal lavage was collected by flushing the peritoneal cavity with 6 ml FACS buffer (PBS with 2 mM EDTA and 0.1% BSA). Mice were gently massaged to ensure optimal collection of peritoneal cells.

For the isolation of thymocytes, the thymi of 3- to 4-week mice were harvested and smashed over a 70-μm strainer to obtain a single-cell solution.

Monocytes were isolated from mouse bone marrow using the EasySep™ Mouse Monocyte Isolation Kit (Catalog #19861, STEMCELL Technologies) according to the manufacturer’s instructions. Briefly, femurs and tibia from 8–12-week-old mice were harvested. Two ends of the bones were cut, and the bone marrow was flushed with a 26 g needle filled with cold sterile 1X PBS through a 70-μm cell strainer to generate a single-cell suspension and subjected to immunomagnetic negative selection. Enriched monocytes were collected as the untouched fraction and used immediately for downstream assays.

### Flow Cytometry

Surface stain antibodies were incubated with cells on ice for 30 min and washed with 1 ml FACS buffer (PBS with 2 mM EDTA and 0.1% BSA). Counting beads (424902; BioLegend) were added to each sample to determine cell numbers. Samples were analyzed on a BD FACSymphony A3 or A5-SE Cell Analyzer using FlowJo software. All antibodies were used at 1:200 dilution.

For intracellular staining, the cells were incubated in Cyto-Fast Fix/Perm buffer (426803, BioLegend) at room temperature for 20 min and then washed with 1× Cyto-Fast Wash buffer (426803, BioLegend). The cells were then incubated with 1× Cyto-Fast Wash buffer (426803, BioLegend) containing cytokine specific antibodies for 20-30 minutes at room temperature. Cells were then washed in FACS buffer and resuspended in FACS buffer for flow cytometry analysis.

For IL-4 and IL-13 staining in T cells, peritoneal exudate cells were incubated overnight in T cell media (RPMI 1640 supplemented with 10%FBS, 1% penicillin/streptomycin, 1% sodium pyruvate, 1% MEM NEAA and 50 μM β-mercaptoethanol) and stimulated with anti-CD3/anti-CD28 (1 ug/mL) or isotype for 24 hours before staining and analysis.

### *In vivo* efferocytosis assay

For 1 hour uptake experiments, apoptotic Jurkat cells were used. Jurkat cells were treated with 50 μM of Hydrogen Peroxide overnight to induce apoptosis. This method typically generates ∼70% annexin V+ cells. The apoptotic cells were resuspended in 1x PBS at a concentration of 1 x 10^7^ cells/mL with 5 mL of Vybrant® DiD cell-labeling solution (Invitrogen) for 10 min. Cells were then washed with PBS before being resuspended in PBS at a density of 1 x 10^7^ cells/mL. Cells were injected with 100 μL of PBS and mice were euthanized 1 hour later and analyzed by flow cytometry.

For serial injections of apoptotic cells, irradiated thymocytes were used. Naïve thymocytes were prepared as described above and were irradiated with 20 Gy of ionizing radiation via SARRP. Cells were then incubated at 37C for 6 hours. Cells were counted and resuspended at a density of 1 x 10^7^ cells/mL in PBS and 100 μL was injected intraperitoneally.

### *In vitro* efferocytosis assay

BMDMs were plated in 24-well plates at a density of 2.5 x 10^5^ cells per well and were incubated with DiD-labelled apoptotic Jurkat cells for 1 hour at a 5:1 AC:macrophage ratio. The wells were rinsed with PBS three times to remove unengulfed apoptotic cells before detached using pre-warmed Cell Stripper (Invitrogen) for 20-25 minutes at room temperature. Cells were then analyzed by flow cytometry.

### Adoptive Transfer

Cells were kept on ice, and all steps were carried out in a laminar flow hood using sterile reagents. When applicable, cells were collected and stained as described above and were sorted using flow cytometry into the indicated populations. Post sort cells were pelleted (500 × *g*, 5 min at 4 °C), resuspended in PBS and counted by Countess (Invitrogen). For monocyte adoptive transfers, 1 x 10^6^ cells per indicated population were injected intraperitoneally. For Tim4-adoptive transfers, 5-10 x 10^4^ cells of the indicated population were transferred. For inflammatory monocyte-macrophage adoptive transfers 2.5 x 10^5^ were injected per mouse into mice previously injected with 1000 μg of Zymosan. For BMDM adoptive transfers 1 x 10^6^ cells were transferred per indicated population. Depletion antibodies targeting CD4+ T cells (150 μg per mouse per dose) were injected on days 7 and 14 after transfer if indicated. Clodronate Liposomes (Liposoma) were used to deplete resident macrophages where indicated as previously described^15^ at 0.0625 mg per mouse in 100 uL 1 week before adoptive transfer.

### Tumor Model

For KPCA tumors, 1 × 10^6^ KCPA cells were injected i.p. into Ms4a3^Cre^; Rosa26^TdTomato^ mice in 200 μL of a 1:1 matrigel:PBS mix (Matrigel Matrix Basement Membrane, 354234; Corning). On day 21 mice were euthanized, peritoneal lavage was collected as described above and omental tumors were collected. Tumors were digested using a cocktail of 1 mg/ml collagenase IV and 100 µg/ml DNase I, in 4 ml DMEM with 10% FBS. The tissues were minced into small pieces prior to digestion at 37°C for 30 min shaking at 800 rpm. Samples were then passed through a 70 µm cell strainer to collect single-cell suspension to be analyzed by flow cytometry. If necessary, single-cell suspensions were treated with 1X red blood cell lysis buffer (555899; BD Bioscience) on ice for 10 min.

### Statistical Analysis

Statistical analyses were performed in Prism (GraphPad Software, Inc.). The specific statistical tests used - Student’s t test, paired t-test, one-way or two-way ANOVA with a post-hoc Tukey’s multiple comparisons test - and other relevant details are noted in the figure legends. Results were considered significant at P < 0.05. Results display all replicated experiments and are presented as mean ± SEM.

### Illustration

Graphical abstract and Figure 3A were generated using https://Biorender.com.

## Acknowledgments

We thank Wistar core facilities (imaging, flow cytometry, and animal facility). This study was supported by NIGMS Maximizing Investigators’ Research Award (R35GM162440; N. Zhang), PA CURE Program (SAP4100095623; N. Zhang), Ovarian Cancer Research Alliance (MIG-2026-1-2197; C. Shao), and NIH Wistar Training Program in Basic Cancer Research (T32 CA009171; N. Param and B. Murphy).

## Author contributions

Conceptualization: B.S. Manning, R. Shinde, M. Carroll, N. Zhang. Investigation: B.S. Manning, L. Zhu, B. Murphy, M.M. Soliman, C. Shao., N.J. Param, T. Miyamoto. Resources: N. Zhang Data Interpretation: B.S. Manning, B. Murphy, M. Carroll, N. Zhang. Writing: B.S. Manning, N. Zhang. Supervision: N. Zhang, M. Carroll. All authors edited the manuscript

---

**Figure S1.**
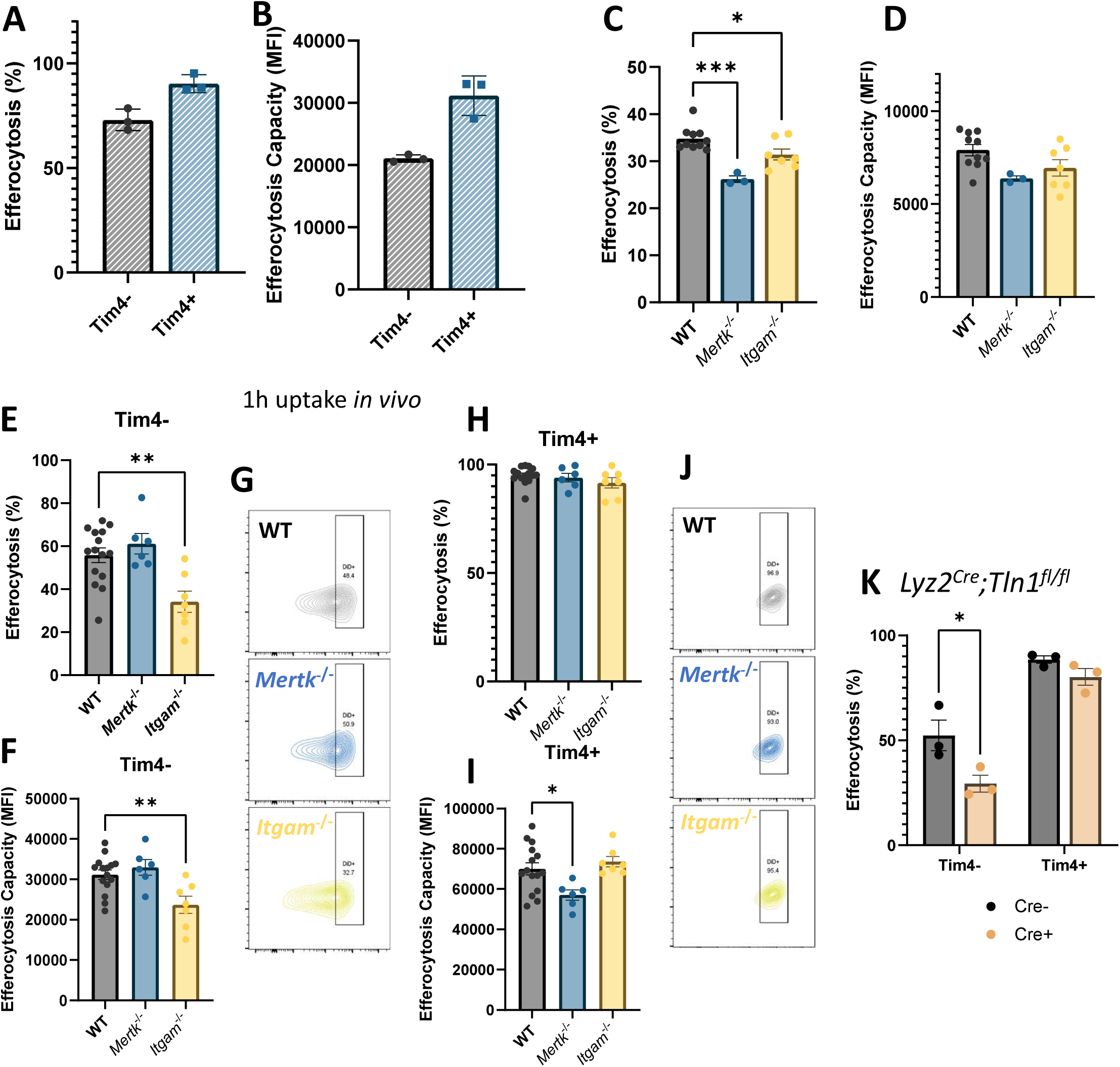
Peritoneal macrophage efferocytosis uses integrin αMβ2. (A-B) In vivo efferocytosis efficiency (DiD+ macrophages) and capacity (DiD mean fluorescence intensity, MFI) in WT macrophages. (C-D) In vitro efferocytosis efficiency (DiD+ macrophages) and capacity (DiD mean fluorescence intensity, MFI) in BMDMs from WT, *Itgam^-/-^* and *Mertk^-/-^* mice (E-G) In vivo efferocytosis efficiency (DiD+ macrophages) and capacity (DiD mean fluorescence intensity, MFI) in Tim4^-^ peritoneal macrophages from WT, *Itgam^-/-^* and *Mertk^-/-^* mice with representative contour plots. (H-J) In vivo efferocytosis efficiency (DiD+ macrophages) and capacity (DiD mean fluorescence intensity, MFI) in Tim4^+^ peritoneal macrophages from WT, *Itgam^-/-^* and *Mertk^-/-^* mice with representative contour plots. (K) In vivo efferocytosis efficiency (DiD+ macrophages) in Tim4+ and Tim4-peritoneal macrophages from *Lyz2^Cre^;Tln1^fl/fl^* and Tln1^fl/fl^ mice

**Figure S2.**
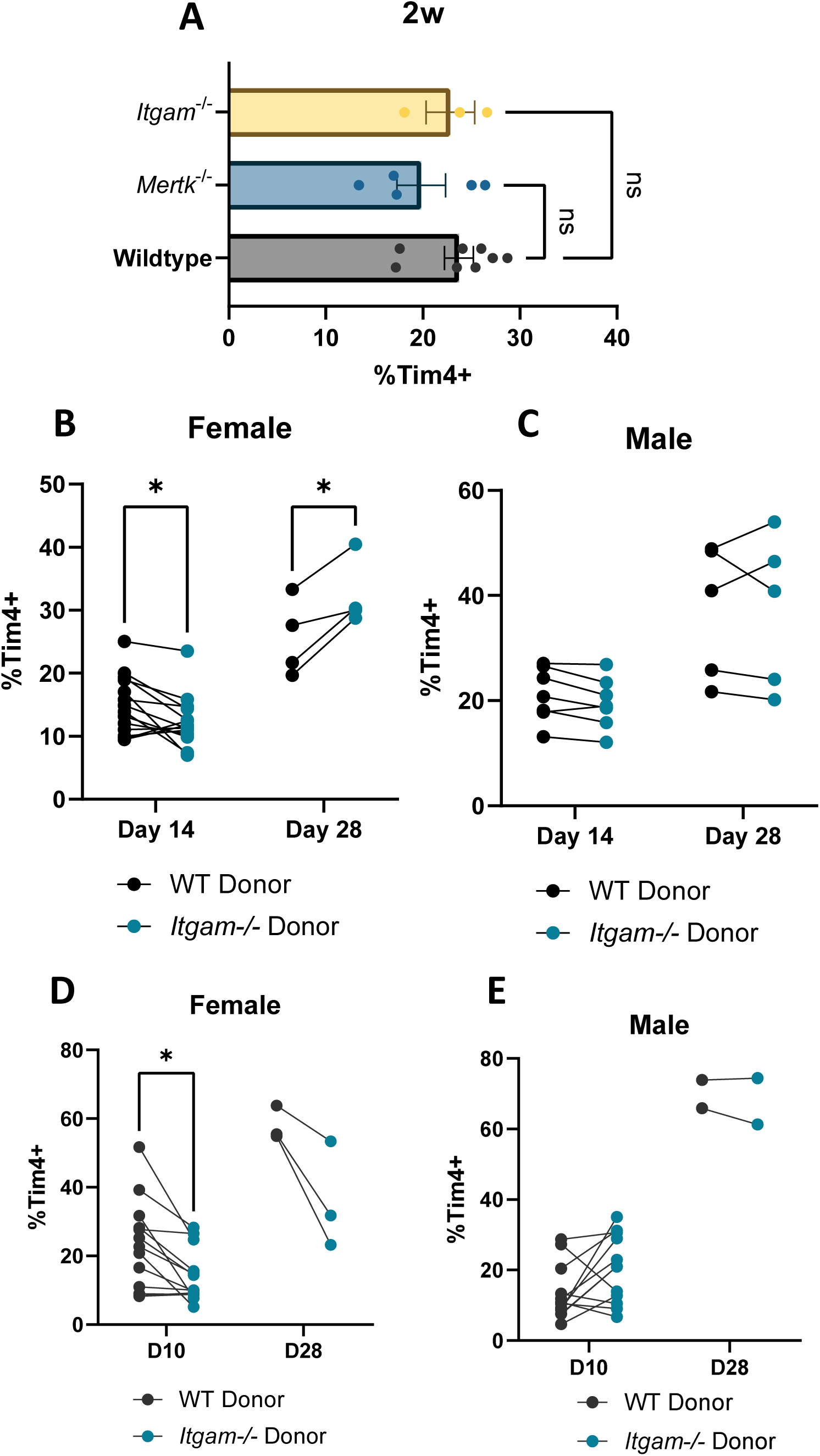
Loss of integrin αMβ2 alone does not consistently impair Tim4+ differentiation. (A) Percentages of Tim4^+^ from CD45.1 WT and CD45.2 *Itgam^-/-^* or *Mertk^-/-^* peritoneal macrophages from mixed bone marrow chimeras 2 weeks after transplant. (B-C) Percentage of donor CD45.1^+^ and CD45.2^+^ *Itgam^-/-^* bone marrow monocytes that differentiated into Tim4^+^ peritoneal macrophages after the indicated time frame in male and female mice. (D-E) Percentage of donor CD45.1^+^ and CD45.2^+^ *Itgam^-/-^* FACS-isolated monocyte-macrophages from mice 72 hours after treatment with 1mg of Zymosan.

**Figure S3.**
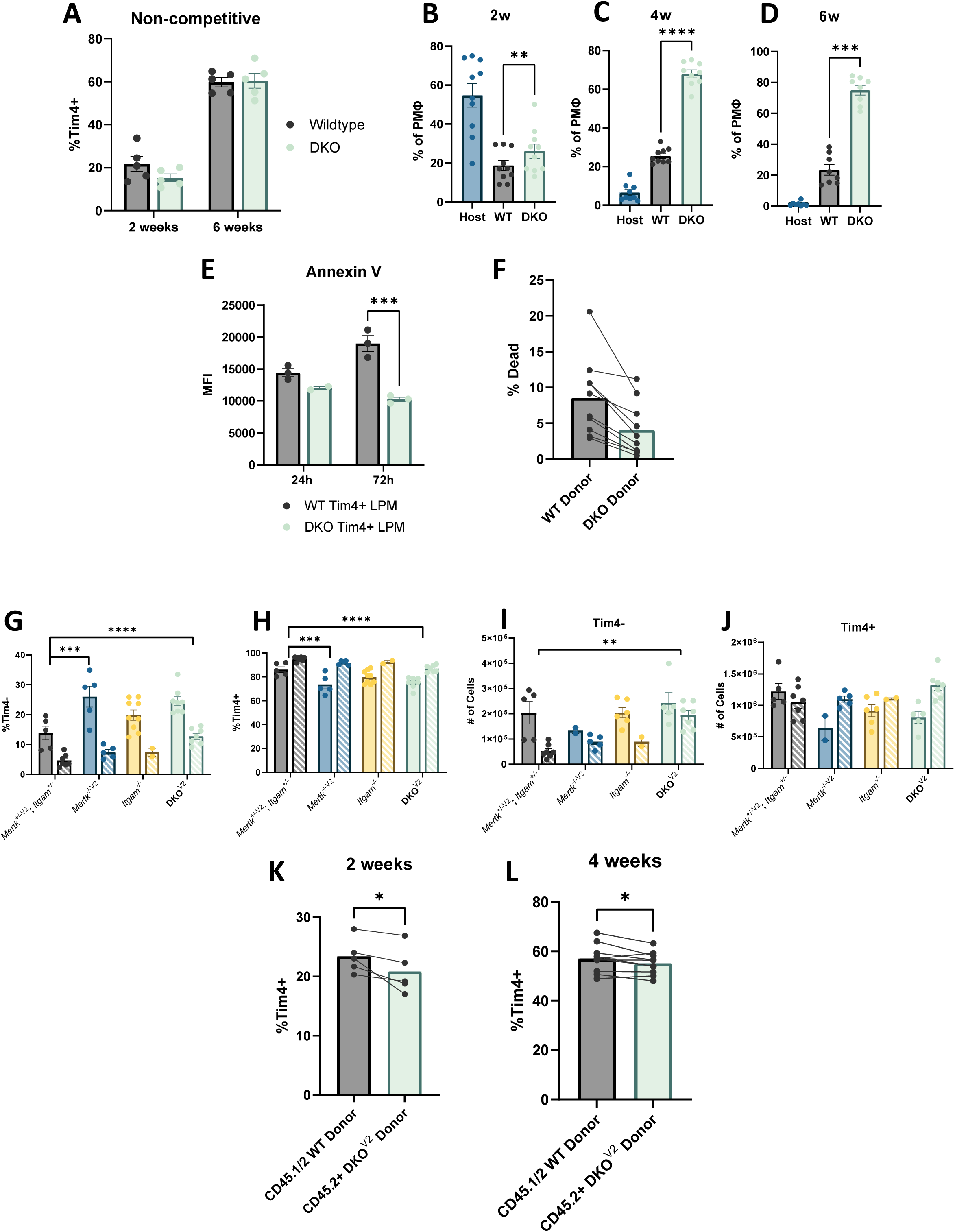
(A) Percentages of Tim4^+^ from CD45.1 WT and CD45.2 DKO peritoneal macrophages from single donor bone marrow transplant 2 and 6 weeks after transplant. (B-D) Composition of peritoneal macrophage pool from donor CD45.1^+^ CD11b^Hi^ WT and CD45.2^+^, CD11b^-^ DKO and host CD45.2^+^, CD11b^Hi^ at 2-, 4- and 6-weeks after transplant. (E) Annexin V MFI of FACS-isolated Tim4^+^ peritoneal macrophages from WT and DKO mice cultured in vitro for 24- and 72-hours. (F) Percentage of Live Dead Yellow+ peritoneal macrophages from CD45.2^+^ WT and CD45.2^+^ DKO cells from adoptive transfer experiments into NSG mice. (G-H) Proportion and (I-J) Quantification of peritoneal macrophages that are Tim4^-^ and Tim4^+^ from Het;Het, Mertk^-/-V2^, Itgam^-/-^, and DKO^V2^ 8–10-week-old mice. Data are presented as mean ± SEM. Statistical analysis was performed using two-way ANOVA with Tukey’s multiple-comparisons test. P values shown indicate significant differences for the main effect of genotype (column factor), independent of sex. *P < 0.05, **P < 0.01, ***P < 0.001, ****P < 0.0001. (K-L) Percentages of Tim4^+^ from CD45.1 WT and CD45.2 DKO^V2^ peritoneal macrophages from mixed bone marrow chimeras. Each point represents an individual mouse. Lines connect paired measurements from the same mouse. Statistical significance was assessed using a paired two-tailed Student’s t-test. *P < 0.05, **P < 0.01.

**Figure S4.**
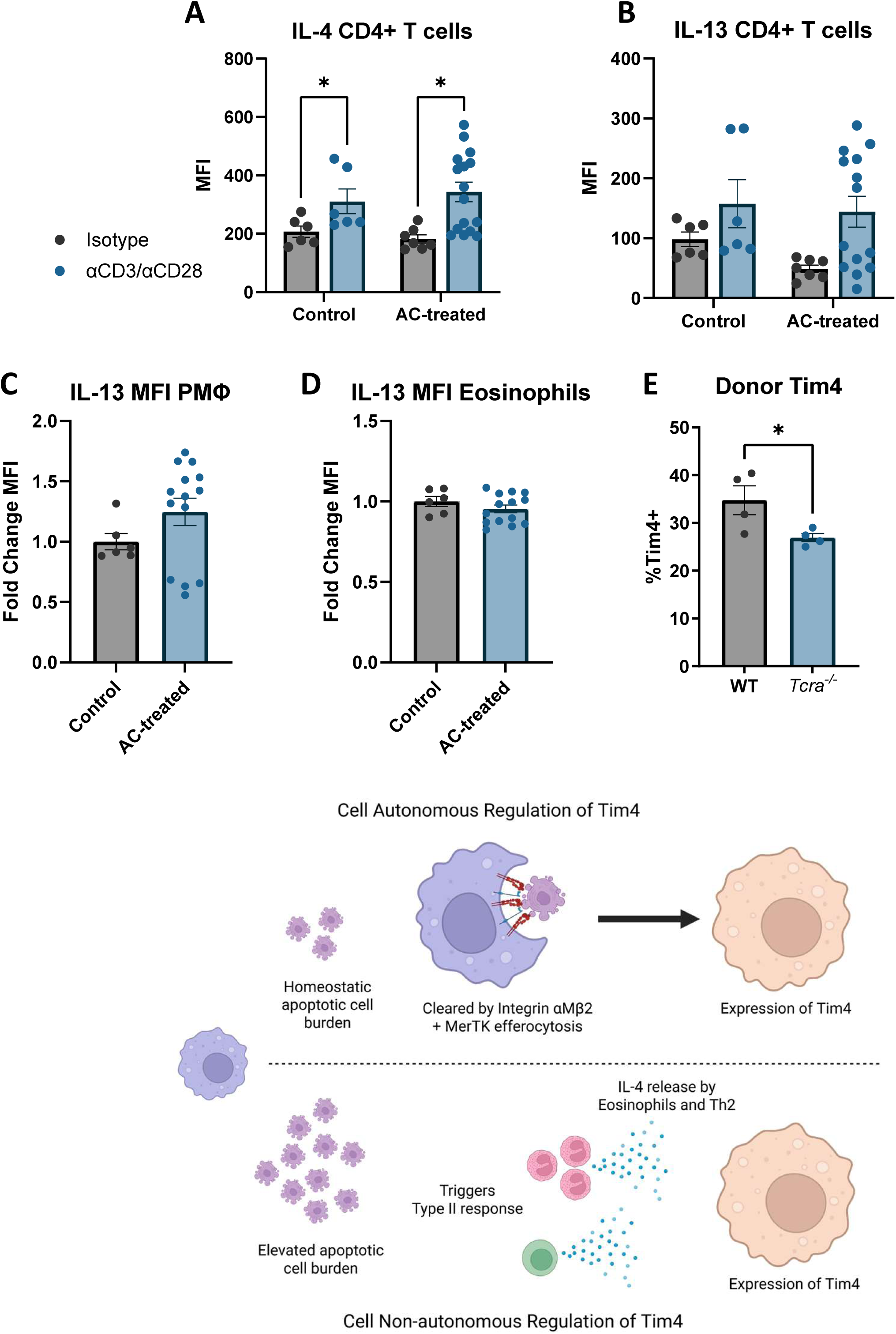
(A-B) Intracellular IL-4 and IL-13 MFI of CD4+ T cells on day 15 of the immune response 24 hours after stimulation with αCD3/αCD28 or isotype. p < 0.05(*), paired student’s t-test. (C-D) Intracellular IL-13 MFI of peritoneal macrophages and eosinophils on day 15 of the immune response. (E) Proportion of Tim4+ donor peritoneal macrophages 18 days after BMDM adoptive transfer into WT or *Tcra^-/-^* mice with apoptotic cell injections.

